# A brain-rhythm based computational framework for semantic context and acoustic signal integration in speech processing

**DOI:** 10.1101/2024.01.17.575994

**Authors:** Olesia Dogonasheva, Keith Doelling, Denis Zakharov, Anne-Lise Giraud, Boris Gutkin

**Affiliations:** Group of Neural Theory, École Normale Supérieure PSL^*^, Paris, France; Institut de l’Audition, Institut Pasteur, Université de Paris Cité, Paris, France; Centre for Cognition and Decision Making, HSE University, Moscow, Russia

**Keywords:** rhythms, predictive coding, speech recognition, inference model, invariant speech processing, auditory cortex

## Abstract

Unraveling the mysteries of how humans effortlessly grasp speech despite diverse environmental challenges has long intrigued researchers in systems and cognitive neuroscience. This study explores the neural intricacies underpinning robust speech comprehension, giving computational mechanistic proof for the hypothesis proposing a pivotal role for rhythmic, predictive top-down contextualization facilitated by the delta rhythm in achieving time-invariant speech processing. Our Brain-Rhythm-based Inference model, BRyBI, integrates three key rhythmic processes – theta-gamma interactions for parsing phoneme sequences, dynamic delta rhythm for inferred prosodic-phrase context, and resilient speech representations. Demonstrating mechanistic proof-of-principle, BRyBI replicates human behavioral experiments, showcasing its ability to handle pitch variations, time-warped speech, interruptions, and silences in non-comprehensible contexts. Intriguingly, the model aligns with human experiments, revealing optimal silence time scales in the theta- and delta-frequency ranges. Comparative analysis with deep neural network language models highlights distinctive performance patterns, emphasizing the unique capabilities of a rhythmic framework. In essence, our study sheds light on the neural underpinnings of speech processing, emphasizing the role of rhythmic brain mechanisms in structured temporal signal processing – an insight that challenges prevailing artificial intelligence paradigms and hints at potential advancements in compact and robust computing architectures.

## 1 Introduction

Speech processing, with its inherent complexities and multidimensional nature, continues to be a focal point of cognitive neuroscience. In part, this is due to its highly robust nature: humans comprehend speech across a wide spectrum of voices, ranging from young children to elderly individuals, from speakers of different languages to regional dialects, and even across diverse socio-cultural backgrounds. Moreover, speech comprehension remains robust despite variations in speech rates, encompassing both rapid and leisurely speech patterns.

A key feature of speech comprehension which may impart this robustness are temporal windows of integration through which sensory information is accumulated and ultimately decoded into speech symbols. These windows are not arbitrarily long but instead are constrained to specific time lengths and at multiple scales [1, 2]. Such a mechanism both allows for evidence accumulation, generating candidate targets while reducing effects of noise, and also focusing on time scales particularly relevant for speech signals. Poeppel (*Speech Communication*, 2003) proposed that the auditory cortex contained populations sensitive to two major time scales: one at ~200 ms, corresponding roughly to the duration of a syllable, and one at ~30 ms, corresponding roughly to the duration of a phoneme [3]. Since this influential proposal, other time scales have also been proposed, in particular on the order of 1 second, as a means of extracting prosodic and phrasal boundaries [4, 5]. Thus, a hierarchy of temporal windows is scaffolded to prioritize the processing of speech relevant units.

Brain rhythms emerge as a compelling candidate for the neural mechanisms supporting these temporal windows of integration in speech processing [4, 6–8]. Substantial empirical evidence indicates that rhythmic brain activity maintains a hierarchical structure during the processing of speech, and this hierarchy aligns with the inherent structure of speech itself [9–11]. It is thus plausible to posit that the rhythmic structure of speech interacts with the scaffold of endogenous brain rhythms, thereby establishing temporal processing windows. These windows, in turn, govern the real-time processing and comprehension of incoming auditory signals [8, 12].

Synchronization of neural activity with a rhythmic stimulus, such as speech [13– 15], might be seen as a natural mechanism for the opening and closing of these windows. In the primary auditory cortex, for instance, the theta rhythm is acknowledged to be entrained by the speech envelope, thereby encoding syllabic information [12, 16–18]. Concurrently, oscillations in the gamma range embedded within a theta-cycle have been shown to encode phonemes [19, 20], giving rise to a theta-gamma code for syllables [7, 21] mapping well to the originally proposed timescales of Poeppel (2003). Previous investigations have proposed that the theta-gamma code orchestrates a bottom-up information flow, starting from sounds captured by the cochlea and converging in the primary auditory cortex [22–24]. This rhythmic windowing, characterized by theta-gamma dynamics, confers robustness to speech parsing in noisy and compressed speech scenarios [24, 25].

The proposal of brain rhythms as a neural mechanism of temporal windowing leads to specific predictions for how the comprehension system should respond to temporal compression and distortions. Speech comprehension should be largely impervious to temporal distortions as long as the neural timescales can still synchronize with those of the input. This has been borne out in multiple studies using temporal interruptions and segmentations. In experiments with interrupted speech [26], silent intervals masked the speech at different time frequencies, i.e., the speech signal was interrupted by silences. As a result, some elements of the speech were simply missing. The results of the experiment showed that when the frequency of the interruptions was greater than 1 Hz (500 ms of signal, 500 ms of silence), speech recognition recovered to nearly control levels.

In another set of studies, silent intervals of different durations (up to 500 ms) were inserted into the speech. Here, the segmented signal contained all the parts of the original speech, and no information was deleted [27]. In these experiments, subjects’ performance showed characteristic U-shaped curves, with the worst performance when the silence durations were 100 ms, whatever the silence-to-speech ratio. In another manipulation, speech was compressed by different factors. Subjects in these tasks showed robust success in recognising speech as long as the compression factor was less than 2, above which performance dropped catastrophically [28–31]. Intriguingly, when this temporally squeezed and incomprehensible speech was split into chunks interspersed with silences, recognition recovered [24, 30, 32–34]. Here, the performance errors showed a characteristic U-shape with the fewest errors when the overall natural duration of speech was restored by the silent insertions. These results underline the importance of aligning the temporal scales of speech with endogenous scales set by multiple brain rhythms in reconstructing meaning from the acoustic speech flow. Understanding the mechanisms that enable humans to navigate through this large parameter space presents an intriguing challenge and, arguably, a litmus test for the potential neural mechanisms underlying speech recognition processes. Notably, how could we explain why speech comprehension is recovered by adding silences that do not carry any information? The most plausible explanation for this phenomenon lies in the hierarchical rhythmic structure inherent in meaningful speech, significantly contributing to robust and temporally invariant comprehension [35–38].

The hierarchical configuration of time scales in speech and neural processing is met with a similar hierarchy in natural language, which is also pivotal for the efficiency of human speech processing [39–43]. Such hierarchical organization spans all linguistic levels, from the phonetic structure of words to the highest tiers of communication [44]. While the hierarchy of neural rhythms can support the opening of temporal windows of the right size for speech decoding, it has not been used to describe the underlying analysis of these linguistic levels and how they interact. This line of reasoning prompts a fundamental inquiry: How does the hierarchy of linguistic representation interact with the time scales of speech perception to generate neural mechanisms that support the effective processing and comprehension of speech?

As a key conceptual proposition in this paper, we suggest that a top-down predictive information flow modulated by delta rhythm can mitigate the deterioration of speech signals and improve processing reliability in acoustically challenging environments [45–50]. More specifically, we hypothesize that the information from multiple syllables is predictively combined into a semantic contextual representation (e.g., a word or a prosodic phrase) via a process indelibly intertwined with the delta rhythm [51]. Nevertheless, the computational mechanisms governing the formation of such predictive representations and how this process distinctly contributes to speech processing in the brain remain open questions.

The delta rhythm, being the slowest rhythm observed in the auditory cortex during speech processing, may be a plausible carrier of contextual top-down predictions [52]. Studies suggest that the functions of the delta rhythm include tracking of prosody [53–57], chunking of words and phrases [5, 58], error resolution [59, 60], multiscale integration [51, 61], top-down modulation of speech processing [46, 62, 63], as well as top-down prediction of temporal information [64, 65]. However, existing models primarily employ the delta rhythm as a mechanism for chunking words and phrases [66, 67], overlooking its potential role in top-down contextual influence.

To show how top-down contextual influences integrate with bottom-up signals, forming resilient and consistent speech representations and processing, we propose the Brain-Rhythm-Based Inference model (BRyBI). BRyBI mechanistically incorporates diverse brain-rhythm data and shows time-invariant speech processing. In this model, hierarchically organized interacting rhythms actively sustain the flow of both top-down and bottom-up information during the inference process: theta-gamma interactions delineate and parse the phoneme/syllable sequences, while the delta rhythm dynamically generates the inferred word/prosodic-phrase context. We demonstrate how these processes facilitate speech recognition even in complex conditions. Additionally, we elucidate the mechanisms underlying the remarkable recovery of comprehension of perturbed speech when specific timescales of the spoken rhythm are re-established. We propose that rhythmic predictive top-down contextualization plays a pivotal role in explaining time-invariant speech processing. Furthermore, our model predicts the restoration of comprehension in compressed speech through the re-chunking of words and phrases, emphasizing the critical dependence on top-down delta-dependent processes.

## 2 Results

### 2.1 Conceptual structure of the rhythm-based Bayesian inference computation for speech processing

Our proposed model is fundamentally rooted in the predictive coding framework, wherein prior information is encoded within an internal model of the environment, often referred to as a generative model (GM) [68, 69]. This internal model actively influences perception [70]. The GM generates predictions of sensory signals, and these predictions are subsequently compared with the actual incoming peripheral signals. The resultant comparison yields prediction errors that traverse the model hierarchy to update the internal states within the GM. Multiple studies have demonstrated that the predictive coding framework provides a plausible paradigm for audio perception [71]. Firstly, predictive coding reproduces a hierarchical structure that emphasizes linguistic organization and the hierarchy of speech processing [43, 72]. Secondly, because states in a predictive model are dynamic BRyBI closely reflects the real nature of brain processes, dynamically integrating top-down predictions and bottom-up mismatch errors, enabling real-time speech parsing. Recent advances in predictive coding models have demonstrated a balanced implementation of linguistic aspects and the mechanistic plausibility of biophysical algorithms for speech processing [23, 73–75]. Consequently, we have implemented the BRyBI model as a predictive coding model, wherein bottom-up and top-down rhythm-based processes are structured along a theta-based code for syllable parsing (bottom-up) and a delta-based top-down predictive code for phrase parsing and comprehension.

The generative model in BRyBI is structured to have two-levels. The bottom and top levels of the BRyBI speech processing hypothetically map onto the primary auditory cortex (pAC) and the associative auditory cortex (aAC) [60], respectively (Fig. 1). At the top level, the delta rhythm provides the temporal scaffold for semantic context influence, while coupled theta and gamma rhythms at the bottom level encode the acoustic signal of speech, depending on the context. The context represents the prosodic phrases and sets predictions for the sequence of the constituent syllables and phonemes.

**Fig. 1.**
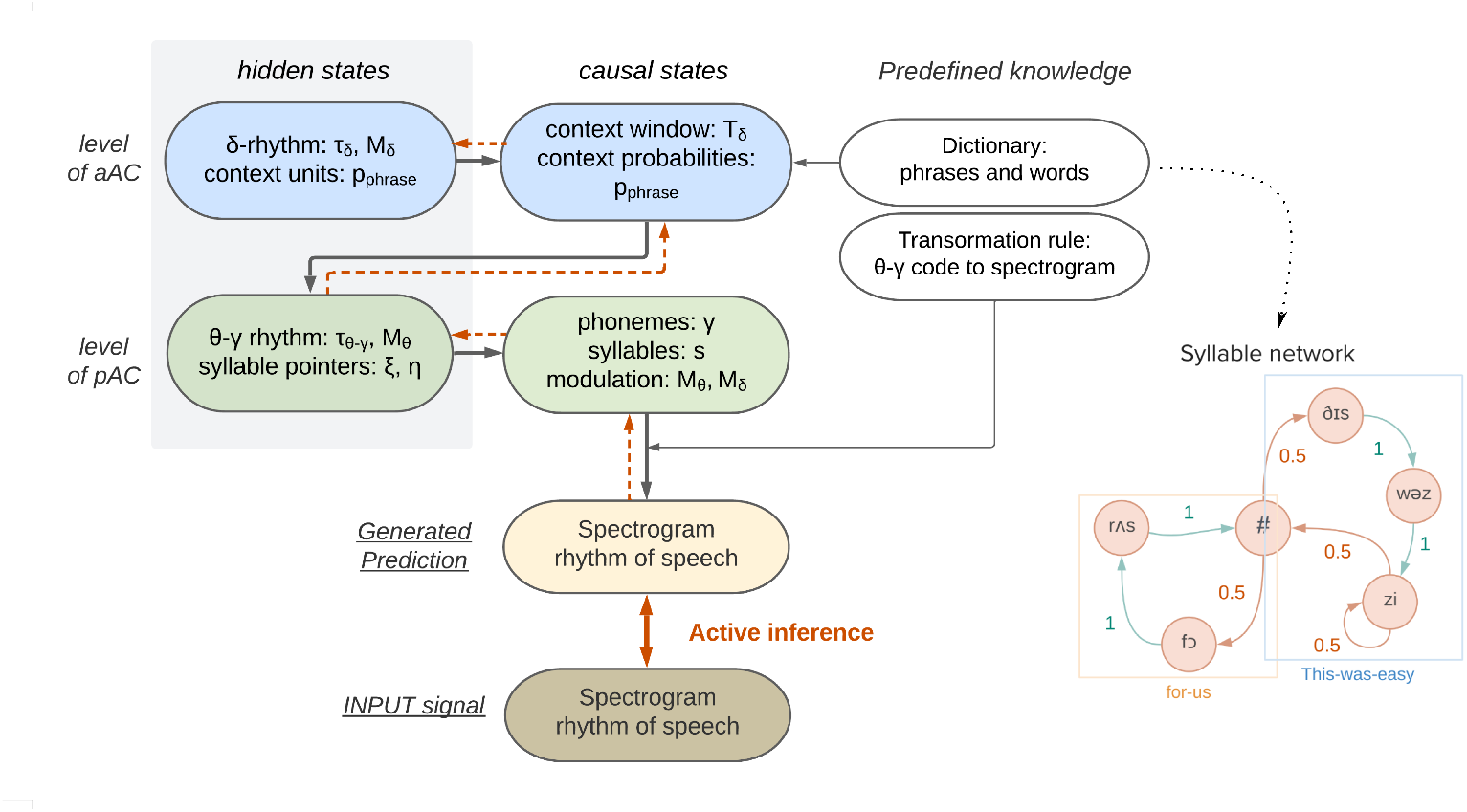
The BRyBI model incorporates predictive Bayesian inference for rhythm-based dynamical speech formation. The hierarchy comprises levels of primary auditory cortex (pAC) and association auditory cortices (aAC). At the aAC level, the delta rhythm governs semantic context formation as an expected prosodic phrase and passes it to pAC level, where coupled theta-gamma rhythms encode the acoustic signal conditioned by the context. The context is represented as a syllable network where each node is a syllable, and connections are possible transitions between syllables. Weights on connections is a confidence in the transition. The sign “#” is a pause that separates words and phrases. The pAC level combines information from both top-down and bottomup flows, creating a theta-code of speech. The theta-gamma code is transformed into a spectrogram through the convolution with predefined tensors (transformation rule). The generated prediction is the spectrogram and speech rhythm (modulation signal at delta rhythm and modulation signal at theta rhythm). The prediction error is passed bottom-up and used to infer the next prediction. Hidden states are represented as dynamical variables (gray background), and causal states are their nonlinear transformation. The generative model supports top-down information flow (black arrows), while the inference provides bottom-up error passing (red dashed arrows) through the hierarchy.

We incorporate a theta-gamma code for syllables in our BRyBI model in accordance with numerous experiments [12, 16, 76–79], where the theta rhythm is shown to be entrained by the speech envelope and thus synchronized with syllables, and the gamma rhythm that is coupled with theta rhythm encodes phonemes. In the BRyBI model, the theta rhythm is also entrained by a rhythm of syllables and performs temporal segmentation of the continuous signal into the syllables. The coupled gamma rhythm is involved in phoneme coding. A similar realization of these mechanisms, where the coupling of theta and gamma rhythms improved speech processing, was proposed [23].

The syllable formation dynamics are rhythmically controlled by theta rhythm in an interactive activation process [80, 81]. The top level of the GM creates a pattern of possible and probable syllable sequence transitions for the current context unit (e.g., word). Figure 1 shows an example syllable network that determines the sequential activation of syllables according to the predicted context unit. Then context is determined as the probability of transitions between syllables in the network.

To gain some intuitive insight, let us consider an illustrative example when the dictionary consists of only two phrases, “This was easy” and “for us” and their constituting words. The input sentence for recognition is “This was easy for us” (Fig. 2A). The graph for the syllable sequence network is in Fig. 1. The first set of phonemes activates the syllable “ðI”. Within our simple dictionary, conditioned on this first syllable, the system has zero uncertainty (100% of confidence) for the transition to the next two syllables. The context dictates the activation of the syllables “sw@” and then “zi” even if the acoustic signal of the phonemes is distorted or is not received. For the subsequent syllable, uncertainty is 50%, because there are two alternatives: to activate the syllable “zi” again or to finish the phrase with a pause denoted by “#”. At this juncture, the system relies on the acoustic signals to reconstruct the spectrogram, rather than generate it based on the contextual flow. The next section describes what this means for delta vs theta stimulus coherence.

**Fig. 2.**
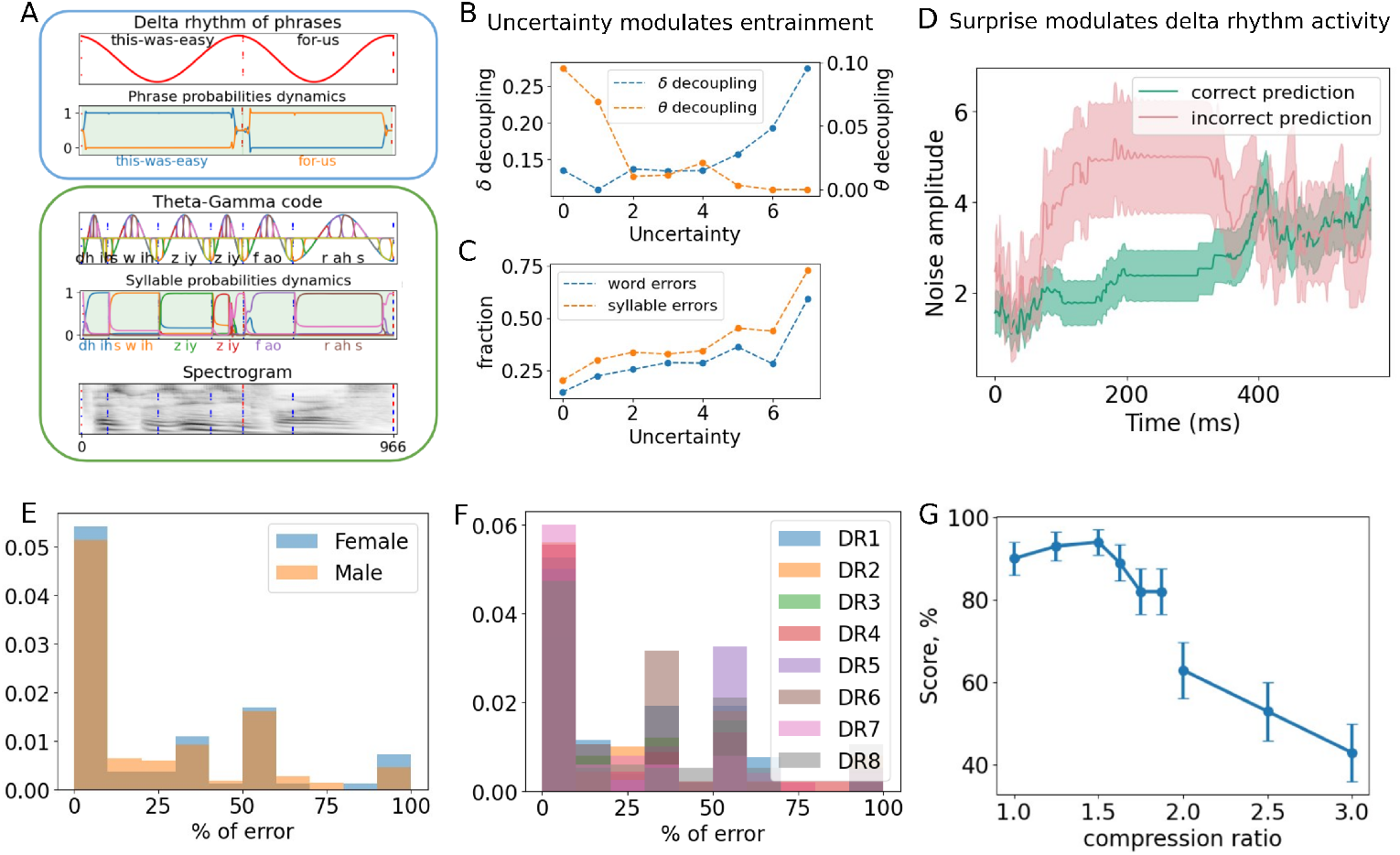
General performance of the BRyBI model for natural speech. (A) An example of speech recognition by BRyBI for the test sentence: “this was easy for us” with two prosodic phrases: “this was easy” and “for us”. During online speech recognition, BRyBI accumulates evidence for the current context unit and infers probability for each phrase at the top level. Similar accumulation happens at the bottom level for syllable probabilities. The green background represents correctly recognized phrases and syllables. Line colors correspond to phrases and syllables (signed on x-axes). (B) Rhythm decoupling depends on uncertainty (represented as numbers of increasing intervals, see Methods). With increased uncertainty, stronger theta coupling with the speech envelope enables reliable phoneme decoding, facilitating speech recognition in unpredictable/noisy conditions. At low uncertainty levels, the context aids predictions (delta-speech coupling relatively pronounced), but theta rhythm locking is not needed. (C) The fraction of errors in words and syllables depends on uncertainty. As uncertainty grows, speech recognition errors increase, though growth is bounded at an uncertainty unit value of 6 due to the theta rhythm locking mechanism. (D) Surprise as Error-Related Potential (ERP) in BRyBI. In cases of incorrect choice of phrase with high confidence (low uncertainty), a prediction error is passed bottom up and causes a change in the chosen phrase. The deviation from the chosen trajectory of dynamic variables occurs at aAC level due to noise addition. The noise amplitude correlates with the deviation and, thus, correlates with an error in the chosen semantic context. (E) Invariant performance in conditions of different speaker genders, and (F) dialects DR1-DR8 in the TIMIT dataset, and (G) speech rates from 1 to 3. When the speech rate is below 2, intelligibility remains consistently high, aligning with an average syllable frequency of less than 10 Hz and an average phrase frequency of less than 4 Hz (Tab. S1). However, a compression ratio exceeding 2 results in syllables and phrases extending beyond the limits of theta and delta rhythms, respectively.

Finally, the system receives the next phoneme, “z”, and can follow a certain context of the phrase “this was easy” with 100% confidence. This example of reconstruction is simplified as much as possible in order to demonstrate the mechanism behind it. In the final stage, GM converts a sequence of syllables into a spectrogram. At each time step, a syllable is selected at the bottom level. Utilizing the gamma rhythm, the syllable is divided into 8 segments (see Methods). Now to compute the spectrogram, we employ convolved pre-trained tensors (see Section Methods) associated with an identified syllable and phoneme. This convolution yields amplitude values for each frequency band associated with the syllable and the phoneme.

To drive the model, we use both the spectrogram and syllable and prosodic envelopes (see details in Methods) as inputs to the bottom level. The original dataset is a preprocessed TIMIT dataset [82]. Using the matlab code [23], we extracted the 6-channel spectra from the sentences as described in Section Methods: Dataset.

Once the model constructs a candidate speech signal segment, the Dynamical Expectation-Maximization (DEM) algorithm is used to infer and optimize the states in the generative model [70]. The states in GM include candidates for current context units (phrases and words), syllables, phonemes, and the phase of delta and theta rhythms (see details in Methods). During the inference, the trajectories in the GM are reconstructed, and inferred phrases and syllables are compared to ground truth phrases and syllables (Fig. 2A).

### 2.2 Rhythm-modulated predictive generative model (BRyBI) recovers speech despite temporal and content perturbations

Speech recognition by the BRyBI model shows good accuracy (15% word error rate for 100 sentences, when the current baseline is 8.3% [83]) for natural speech input (see Fig. 2A for an example). The top-down process plays a crucial role selecting subsequent syllables and phonemes in the model. When context is poorly established, predicting the next syllable becomes challenging, requiring increased sensitivity in theta-syllable synchronization [60, 84]. Such interplay between rhythms is possible through the predictive coding framework. Figures 2B and 2C demonstrate the model’s performance in speech recognition and rhythm entrainment, depending on the uncertainty of the next phoneme. For higher uncertainty, the theta rhythm follows a syllabic rhythm precisely, enabling robust speech perception. As the uncertainty of the future phoneme increases, the speech recognition error increases (Fig. 2C).

This effect can be explained by the specifics of the inference process in the model. The trajectories of all variables are reconstructed by selecting noise in the equations (1). If the uncertainty is large (the context is not chosen), then the trajectories at the upper level specify uniformly distributed probabilities for all transitions between all syllables. In other words, transitions between all syllables are equally likely. DEM minimizes the input-output difference as well as the noise, yet following the prosodic rhythm at the top level of the model does not add new information in the case of high uncertainty. Therefore, in this case, the delta rhythm is less synchronized with the cadence of the phrases.

The opposite situation occurs at the syllable level when uncertainty is high. Syllables and phonemes do not follow any strict contextual unit. They are guessed due to the fitting of the spectrogram into which they are transformed. In this case, it is important to accurately map a syllable and phonemes to each moment in time (for convolution with the desired tensor). Therefore, we observe stronger synchronization between the theta oscillations and the syllable rhythm during times of high uncertainty. When the uncertainty is low, that is, when the activation sequence of the syllables is known, then the generation of the spectrogram occurs mainly due to the context. Hence, the DEM does not optimize the spectrogram by careful fitting of the theta rhythm. Instead, we observe synchronization of the delta rhythm and the cadence of the phrases.

As we can note in Figure 2C, performance error is largely invariant and does not increase until uncertainty reaches a high value (6 a.u.). This performance level invariance is supported by the theta rhythm that locks stronger and stronger to the speech envelope as uncertainty increases. In other words, when the uncertainty of the next phoneme is small (e.g., 0-2 a.u.), the context is easily formed and used for prediction of the next syllable. In this case, the theta rhythm locking is weak (Fig. 2B). This is consistent with the experimental observations in [45, 61] and provides a mechanistic interpretation for these results. As uncertainty increases, in order to preserve syllable prediction quality, phonemes from the acoustic spectrogram need to be decoded more reliably (the bottom-up information flow needs to be prioritized). Hence, the theta coupling with the speech envelope becomes stronger, enabling the resilience of speech recognition in conditions of high uncertainty.

An erroneous context prediction leads to a mismatch between perception and the prediction process. To rectify the error and select a new context in the model, the information about the mismatch is relayed back up the hierarchy. The error is detected as a surprise, defined as an inconsistency between predicted context and observation that leads to increased activity at the top level (Fig. 2D). This causes the noise amplitude at the top level to increase in order to drive context switching. We equate this activity increase at the top of the model hierarchy with the error-related potentials in associative auditory cortical activity [45, 59].

Our numerical experiments show that BRyBI is largely invariant to the speaker’s voice characteristics, e.g., due to speaker gender or dialect (Fig. 2E,F). The model performance follows observed data in recognition scores with speech rates [28, 29]; we see a relative robustness to speech rate until a critical compression ratio, beyond which performance degrades linearly (Fig. 2G). Performance is hypothesized to drop because accelerating speech faster than twice leads to a critical reduction in the length of the processing windows for syllables [24, 29] (see Table S1 for syllable and phrase duration statistics). Under this hypothesis, syllables that alternate faster than the theta rhythm cannot be parsed. This hypothesis has been tested by examining the limitations of human speech perception of interrupted as well as compressed and repackaged speech [24, 26, 27]. We thus set out to expose our model to such modulated speech signals to examine how rhythm-modulation of the generative inference process may account for the limitations of human perception.

To examine for which human speech recognition patterns the delta-modulated context inference may be a necessary mechanism (congruent with brain mechanisms), we subjected BRyBI to two ablations: a context-free model (Fig. 3A) without the top layer and an arbitrary-timed context model (Fig. 3B) where the top-layer was not rhythmically modulated. We also compared the BRyBI performance with several Large Language Models (LMM) such as the Whisper (OpenAI) model [85], Azure [86], Google Speech-to-text [87], etc.

**Fig. 3.**
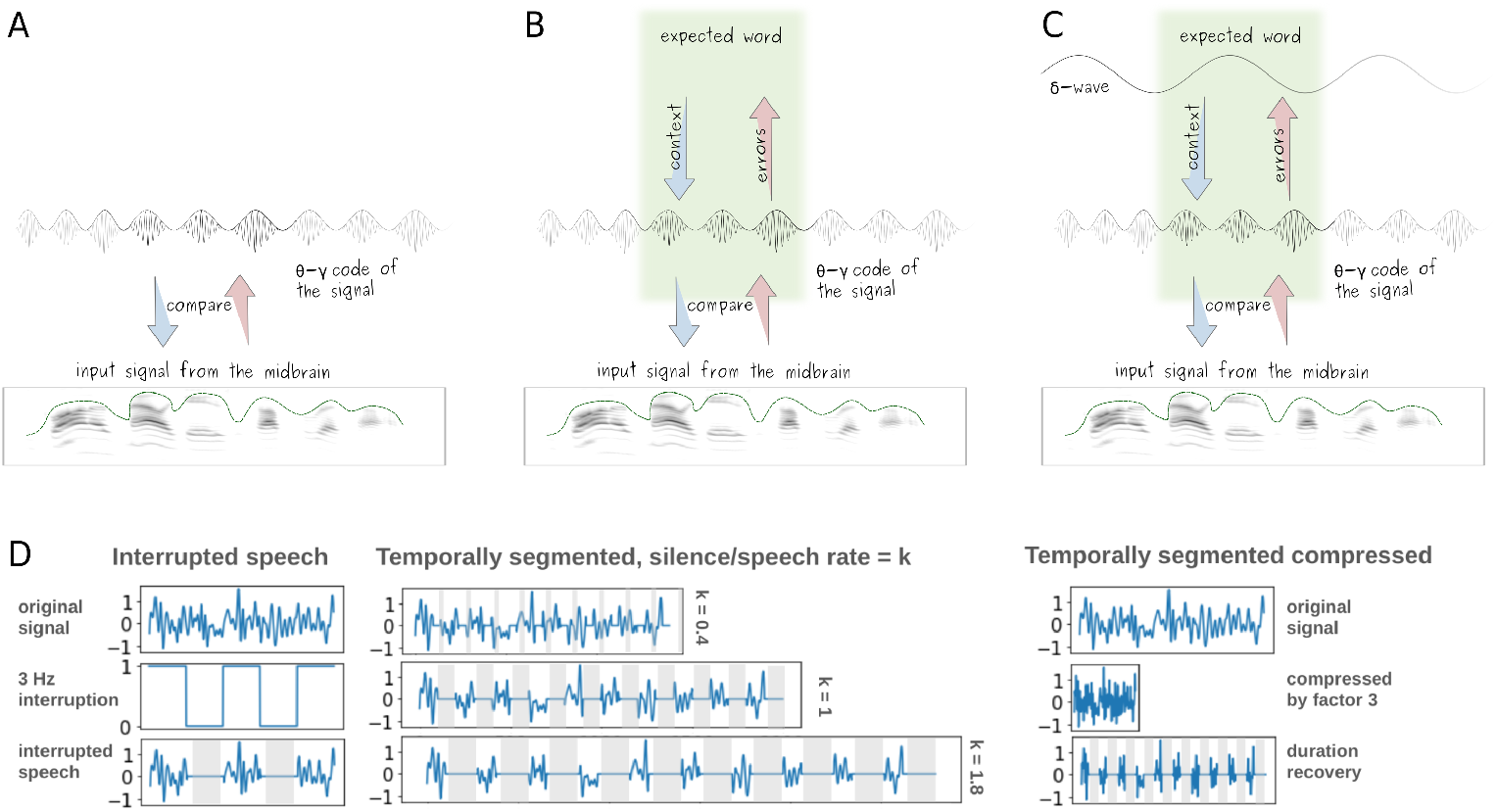
Design of the verification: alternatives for the BRyBI model and experiments with temporary manipulation of speech. (A) The context-free model; (B) arbitrary context model; (C) The dynamic context model (the full BRyBI model). (D) Signal processing for the experiment with interrupted speech is a convolution of the original signal with a rectangular signal. Signal processing for temporally segmented speech includes silence insertion with different silence to speech rate.

We wanted to see which model variants and which LLMs could reproduce specific patterns of error that human subjects make under speech distortions that have been previously observed in experiments. Hence, first, we stimulated alternative/degraded models with speech that is interrupted by silent deletions [26]. Here, following the experiment from [26], normal-speed speech was cut in by a silent interval of various durations, during which speech information was lost (Fig. 3D, left).

The arbitrary and the rhythm-modulated dynamic context versions of BRyBI (Fig. 3B,C) successfully account for the general human behavioral patterns in tasks with interrupted speech (Fig. 4). As in the experiment, contextual BRyBI models exhibit a low performance in the articulation score at the interruption frequency of 1 Hz, which hypothetically can be caused by a long information loss interval (Fig. 4, Fig. S3). We also saw peaks in performance between 10 and 100 Hz, which is the optimal interruption rate where context can recover missed information (Fig. 4, Fig. S3). Finally, the drop at the interruption frequency higher than 100 Hz is provoked by an impairment of phoneme decoding from a distorted spectrogram (Fig. 4, Fig. S3). The context-free BRyBI version showed the performance far from human behavior emphasizing the necessity of contextual support in case of deficient information in acoustic signal.

**Fig. 4.**
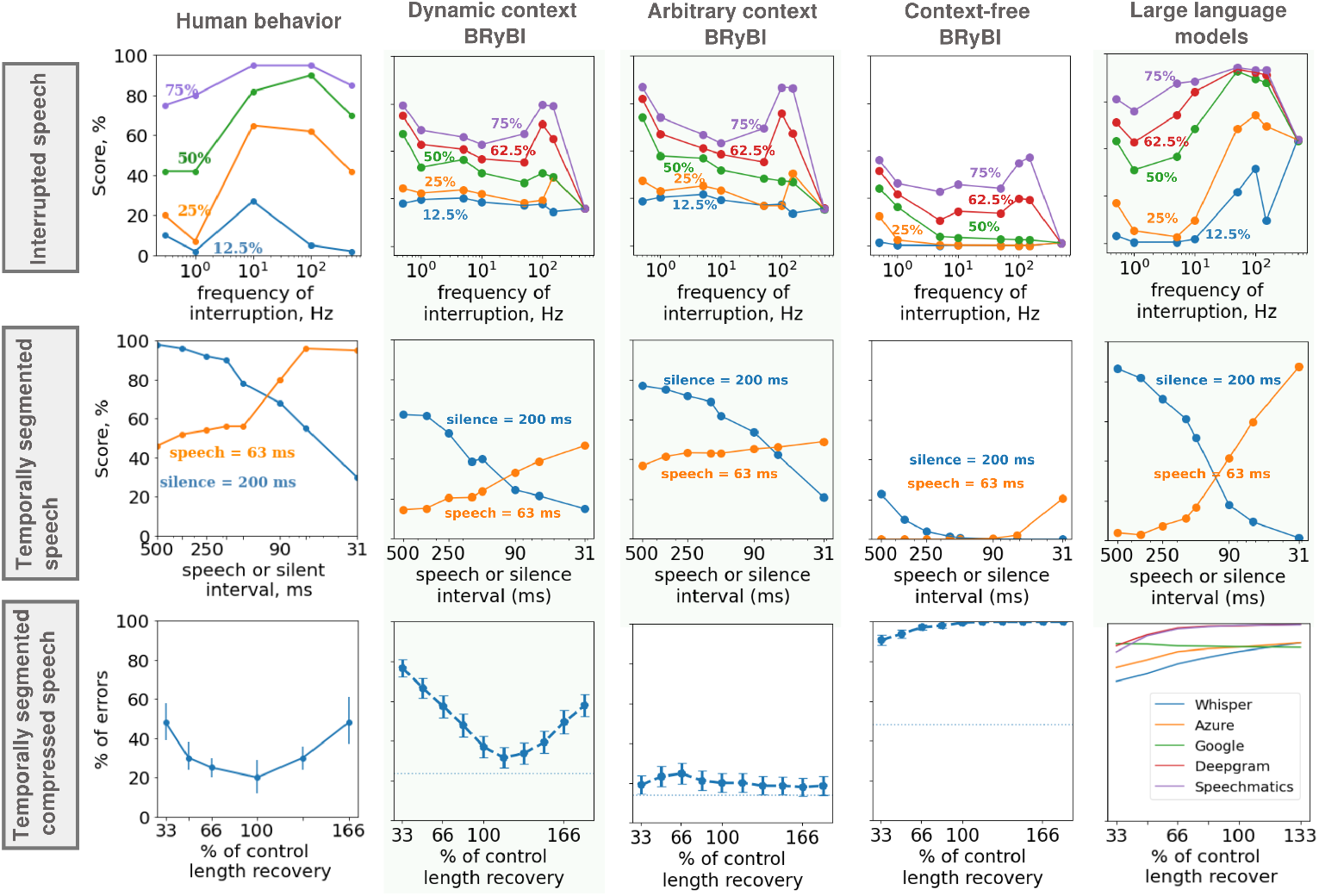
Human behavior and performance of models in experiments with speech under temporal manipulations. The top panel shows results for the experiment with interrupted speech. The middle panel shows results for the experiment with temporally segmented speech. The bottom panel shows results for the experiment with temporally segmented compressed speech. Context-free BRyBI, Arbitrary context BRyBI and LLM do not reproduce the U-shaped curve in the *Ghitza and Grinberg 2009* experiment [24]. This experiment distinguishes the dynamic BRyBI model emphasizing an importance of the context effects which are dynamically governed by delta rhythm. Segmentation of speech by silent intervals does not enhance comprehension for any of the AI models. LLMs we tested do not produce the same pattern of errors as humans, possibly because they are trained for next word prediction and not on silences. The marked difference between LLMs and human speech recognition suggests that the critical missing element in LMMs is the temporal structuring that BRyBI implements utilizing brain rhythms. The behavior data is reproduced from [24, 26, 27].

We then examined how models would perform under speech that was temporally segmented [27]. Here, speech was not interrupted by silent cut-out gaps, but interspaced by added silent segments (Fig. 3D, middle). Thus, there was no loss of information but a change in the timing of segments’ presentation. Maintaining silent intervals at a consistent 200 msec while varying speech intervals reveals a decline in intelligibility for speech intervals ranging from 200 to 31 msec (Fig. 4, second row, human behavior, blue line). Similarly, when speech-interval duration is fixed at approximately 63 msec, increasing silent-interval duration from 63 to 500 msec results in a decrease in intelligibility (Fig. 4, second row, orange line). All models exhibit a behavior coherent with the human behavior for such temporally segmented speech (Fig. 4, second row). Namely, models showed the cross-shaped plot as for speech comprehension, depending on the duration of speech and silent intervals (Fig. 4, Fig. S3). The experiment underscores that the intelligibility of temporally segmented speech depends on the combined durations of speech and silent intervals.

We next turned to a key experiment that we reasoned would allow us to test our main mechanistic hypothesis: that the pattern of invariant speech recovery seen in humans is critically dependent on the delta-modulated top-down inference of the semantic context. In this experiment, speech was modulated by a combination of speech compression and temporal segmentation [24] (Fig. 3D, right). The duration of the silences inserted between segments recovered the natural duration of speech from 33% to 166%. The resulting errors showed a characteristic U-shaped plot of errors in speech recognition depending on the insertion pattern (Fig. 4, human behavior, third row). Neither the context-free model (Fig. 3A) that only implements the bottom level, theta-gamma syllable code, nor the arbitrary-timed context model, where top-level predictions can switch at any time, reproduced the experimental results (Fig. 4, third row). Interestingly, while the arbitrary-timed context model (Fig. 3B) does not produce the telltale U-shape performance, it clearly demonstrates the benefits of context support since it yields low errors across all experimental conditions. The experimentally observed U-shaped error dependency on silence duration is reproduced by adding the delta-band temporal windowing for context alteration in the full model depicted in the Figure 3C.

In order to further show the key role of rhythmically modulated active interference in neural speech processing for the experiment with temporally segmented compressed speech, we compare the BRyBI performance with several LLMs for speech recognition where, to the best of our knowledge, such temporal processes are absent: Whisper (OpenAI), Speech to text (Microsoft Azure), Speech to text (Google), Deepgram, and Speechmatics. These models do not include any notion of temporal windowing for the prediction process. In addition, their models contain vastly more parameters and training points by several orders of magnitude and require much more computational power than BRyBI. All models show low error rates in word recognition for natural speech (2.9 *−* 5.7%, Table S1) and significant degradation for compressed speech over two times (67.9*−*99.1%, Table S1). Furthermore, they perform similarly to human performance in both Interrupted and Temporally segmented speech. However, the models clearly differentiate themselves in the Temporally segmented compressed speech: unlike human behavior and the BRyBI model, repackaging the syllables by silences had no improvement effect on speech comprehension (Fig. 4, third row). From these simulations, we may speculate that even though LLMs show high performance for natural speech, the mechanisms for invariant speech recognition in LLMs and the human brain differ. A key missing element is the temporal structuring of the predictive coding of speech information by endogenous brain rhythms, as it orchestrates the on-line timing of information exchange at multiple scales. We note, however, that the goal of LMM is not to reproduce human behavior but to accomplish language recognition. These results are nevertheless important in the context of recent studies showing a good to almost perfect (up to 100% of the variance) mapping between AI language networks and brain responses [88].

We next examined what the BRyBI model would predict for speech recognition when compressed speech is rechunked by words and/or prosodic phrases. We note that an experiment like this has not been done before. Yet in a comparable study [5] participants listened to spoken digit sequences compressed by a factor of 3 and had to identify a target subsequence. Subjects had best performance when the compressed signals were rechunked by silences bordering the target subsequence boundaries, and that aligned the speech cadence with the delta rhythm. Figure 5 shows an example of the full BRyBI model performance for the experiment with temporally segmented compressed speech. In the control sentence, where no preprocessing was applied, BRyBI correctly reconstructs speech (15% of word errors for 100 sentences). For compressed speech, the model parallels a drop in human behavioral performance. Here, speech compression drives the frequency of phrases in the sentence beyond the delta range. This in turn prevents the model from providing the correct context to help with speech parsing (55% of word errors for 100 sentences). Repackaging the syllables with silences recovers the speech rhythm and thus leads to an improvement in speech intelligibility (Fig. 4, Fig. 5), reproducing the experimental U-shaped curve for repackaging syllables, and predicting the U-shaped curve for repackaging phrases.

**Fig. 5.**
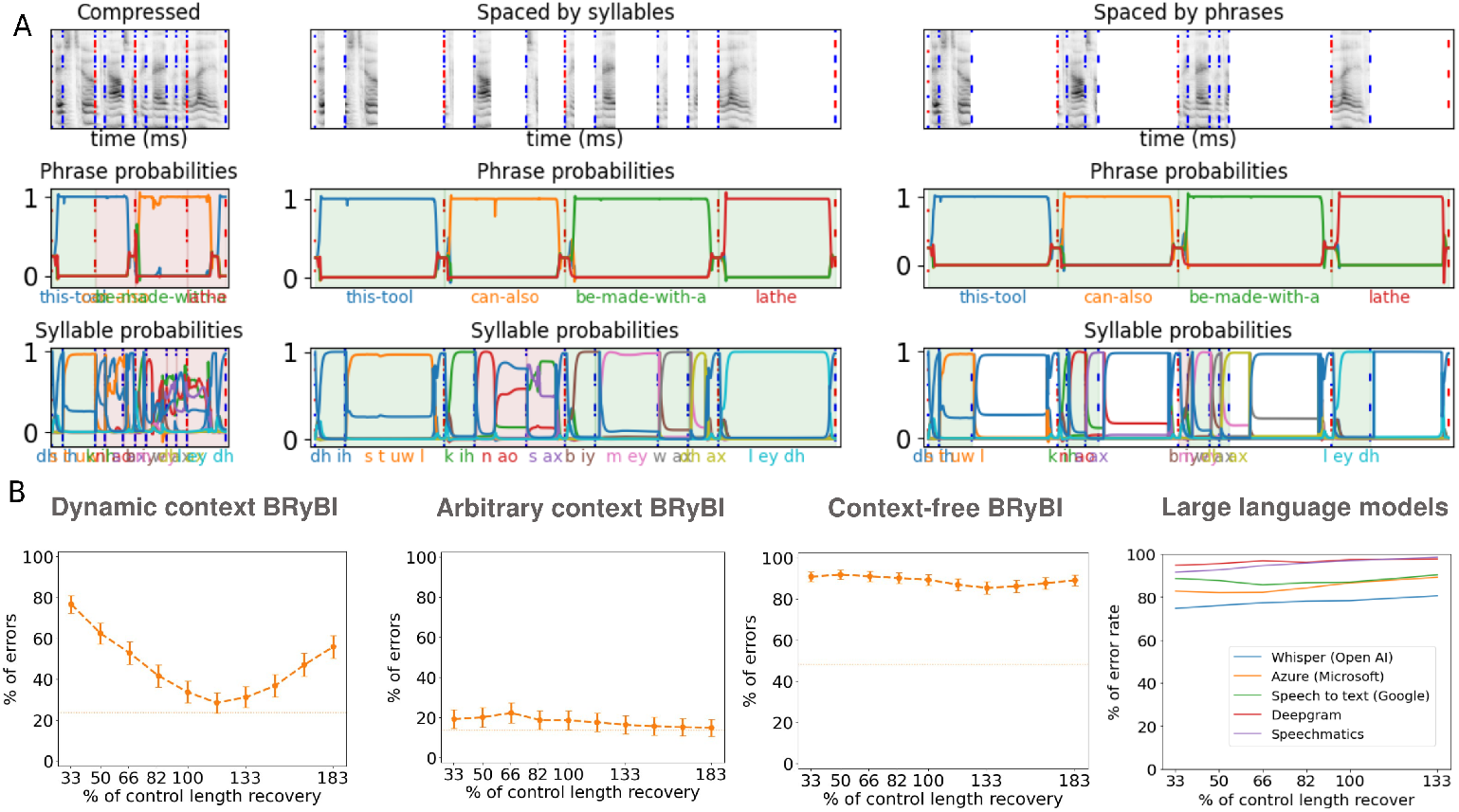
Prediction of the model for compressed speech spaced by prosodic phrases. (A) Example of speech recognition in a test sentence: “this tool can also be made with a lathe”. Correct/incorrect recognition is indicated by the green/red background, respectively. Top panel: spectrogram; middle: reconstructed contextual candidates; bottom: reconstructed syllables; Note the errors in syllable and word sequencing. “Spaced by syllables” for inter-syllable 100 ms silence inserts; “Spaced by prosodic phrases” for inter-phrase 300 ms silence inserts. Compression shortens the time window for context formation and alternation, thus causing errors. Repackaging syllables and prosodic phrases by inserting silent intervals restores speech rhythm and improves intelligibility. (B) Speech recognition by BRyBI versions and LMMs for temporally segmented compressed speech when segments are prosodic phrases. Simulations of dynamic context BRyBI predict the same error pattern when silent intervals are inserted between prosodic phrases.

## 3 Discussion

In this work, we provide a computational framework for understanding the role of brain rhythms in predictive coding, which highlights the importance of the temporal patterning of contextual information and uncertainty in this process. We show how an inferential theta-gamma code, together with the descending predictive influence of delta rhythm, converse in a predictive generative inference model to produce precise and efficient neural processing of speech. The BRyBI model is able to address a threefold challenge. First,it can match patterns of robust and invariant speech processing that are seen in human experiments. Second, it does so in a biologically plausible manner, incorporating several mechanisms crucial to audio processing during speech perception, notably a mechanistically plausible hierarchical structure for syntax processing within a predictive coding structure [35, 42] that is temporally controlled with oscillatory activity [7]. We note that predictive coding has recently received substantial support as a plausible biological framework for speech processing [60, 71]. Finally, by modeling the rhythm generation and content formation phenomenologically, BRyBI achieves its singular performances with a relatively low model complexity, despite its robustness to extraneous speech perturbations. This low model complexity contrasts with the high complexity of the prevalent AI models of speech recognition (e.g., 74M parameters for Whisper AI) and their need for large-scale computing resources, training data requirements, and energy consumption [89]. We note that despite the phenomenological form of the model equations, the inherent model structure emphasizes the biological plausibility of its component processes (rhythms, representation formation, theta-gamma encoding of syllables).

A compelling hypothesis posits that the core mechanism underpinning speech processing within the auditory cortex involves resolving a two-component optimization task within the framework of predictive coding – minimization of uncertainty and surprise [45, 60, 90]. As speech signals are characterized by the dynamic and predictable nature of phoneme transitions, uncertainty here reflects the variable confidence in predicting the next phoneme and surprise denotes the neural response to an unexpected phoneme in the input. Recently, a study combining non-invasive imaging and computational modeling with deep neural networks [60] demonstrated that the word uncertainty reduction can be explained by surprise (thus, updates in GM) and correlates with delta rhythm in aAC, while phoneme uncertainty correlates with modulation of theta rhythm in pAC. The BRyBI model shows how this hypothesis can be substantiated computationally through a synergy of predictive coding and oscillatory activity with only a minimum number of layers. Replicating the processes occurring between the midbrain and the pAC, the bottom level is designed to receive sensory input information, including spectrograms and modulation signals. Using this bottom-up information, the model disentangles phonemes and syllables by minimizing phonemic uncertainty in a way that is compatible with several recently proposed feedforward models [22–24,66]. In BRyBI, a delta-modulated top-down semantic context inference process further guides this bottom-up information flow.

The present computational work shows that top-down context inference and its governance by the delta-rhythm are critical to account for the patterns of human speech processing. In fact, when considering only feed-forward processes, the human performance patterns cannot be reproduced (i.e., the purely feed-forward context-free model fails (Fig.4)). The experiments that we address specifically tested for invariance to degradation, segmentation, and the recovery of performance under re-spacing of audio signals. Critically, the model where the descending context-formation process is not regulated by a brain rhythm, does not reproduce human-like performance; this degraded model is insensitive to speech manipulations. On the other hand, the rhythmically-governed context formation model accounted for both the patterns of invariance and the recovery of distorted speech.

The dynamic context BRyBI model allows us to go beyond just reproducing the phenomenology of behavior but to understand in detail how the predictive coding computations combine with oscillatory temporal governance to orchestrate the necessary brain computations. If we track the computational process within our model, we see that at each time step, the feed-forward module reproduces the dynamics of syllables predetermined for each phrase in the correct order, as captured and governed by the delta module. When a critical discrepancy arises between the generated theta-gamma code and the incoming sensory input, the encoding process at the bottom level deviates from the predicted context. This conflict, in turn, triggers an update of contextual beliefs at the top level. This update induces a sharp shift in the dynamic state of the context module, requiring a transient increase in this module activity (see details in the section: Methods). In essence, consistent with previous findings [60, 91], the reaction to surprise increases delta rhythm activity. Interestingly, this increase in corrective activity reproduces the phenomenology of the ERP signal [45, 50, 92]. We may speculate that the increase in ERP-like neural activity is a signature of updates in generative model states.

Specifically, minimizing surprise online during speech recognition fulfills the goal of selecting the most appropriate context. The judicious choice of context allows the theta rhythm not to be perfectly synchronized with speech, according to the energy minimization hypothesis [45, 61]. Should the context be erroneous, the model effectively needs to update its corresponding state (i.e., the context of a phrase or word). Such a switch requires a time-locked increase in activity at the context level. We can infer that such an increase underpins the error-related potentials seen during complex speech recognition tasks [59, 74]. As a result, the BRyBI model lends support to the hypothesis that top-down predictive and bottom-up acoustic flows are dynamically integrated, as proposed by several studies [23, 49, 93, 94].

Recent studies show a structural hierarchy in the processing of speech features and highlight the relationship between this hierarchy and the organization of rhythmic activity, e.g., [95]. In particular, these experiments showed that the theta rhythm entrainment is correlated with speech clarity and acoustic properties, whereas the delta rhythm is correlated with higher-order speech comprehension. In line with these findings, BRyBI implements processing syntactic speech units in a hierarchical manner, with phonemes and syllables processed at the bottom level and words and phrases processed at the top level. In particular, phonemes and syllables are associated with the gamma and theta rhythms, respectively, whereas words and phrases are associated with the slow delta rhythm. Within BRyBI, the syllable level integrates information from both the bottom-up acoustic signal and the top-down contextual signal.

One of the strongest arguments that questioned the rhythm-based speech parsing is that theta-locking can vary significantly across different experiments. For example, [12] and [29] showed that theta was strongly locked, while other experiments found weak locking despite good behavior performance [55, 96]. We can propose an explanation for this ambiguous evidence using rhythm-based predictive coding for speech recognition. According to our model and previous ones [22, 97], theta-locking is flexible: in clear contexts, it is floating; in unclear contexts, the speech envelope needs to entrain the theta rhythm.

As previously proposed [24], the insertion of silence intervals between chunks of highly compressed speech could restore speech perception by restoring the syllabic rhythm. This hypothesis states that the speed of syllabic information processing is limited by the theta rhythm. This would be due to the fact that phonemic encoding needs to be articulated on-line with syllable decoding by the next hierarchical stage. The speech intake capacity would thus not be limited by phonemic encoding per se, but rather by its readout via descending pathways. The observations when inserting silent gaps between syllable-equivalent chunks do not necessarily support a syllabic-level decoding hypothesis because speech perception may be recovered due to the restoration of a phrasal rhythm rather than the syllabic one. Inserting silences between syllable chunks does not only recover the syllabic rhythm but also the rhythm of words and phrases. At this point, the model results deviate from the previous hypothesis and predict that restoring the rhythm of phrases (even without restoring that of syllables, when silence is inserted only between phrases) could enable speech perception restoration. The following experiment supports this point, and Figure 5 shows that phrase recognition errors decrease when the natural rhythm of the prosody is restored (Fig. 5B). BRyBI simulations thus suggest that the delta rhythm predictively sets the temporal boundaries for speech integration, beyond which speech becomes illegible. Our model results predict that restoring the rhythm of phrases (even without restoring that of syllables when silence is inserted only between phrases) could enable speech perception restoration.

While BRyBI shows promising results, it leads to multiple avenues for extensions and improvements through the implementation of more biological mechanisms for rhythm generation, the incorporation of phase-amplitude coupling (PAC) mechanisms, and considering the role of beta in the inference hierarchy [98].

Another future direction can be to expand and improve the linguistic foundation of the model. For example, several models that propose the incorporation of compositional mechanisms [90, 99–102] can extend the BRyBI model for the semantic part. These models offer an effective and natural implementation of linguistic structures under the framework of interactive-activation models (IAM). The BRyBI model contains IAM as a mechanism of syllable activations. Refinement of this part of BRyBI, following the example of the developments of previous linguistic models, will help to take BRyBI to another level of linguistic plausability. On the other hand, some of these IAMs [90, 99] illustrate language representations processing using asynchrony and inhibition. A biophysical version of BRyBI, e.g., where rhythms are implemented with dynamical neural mass- or spiking-networks [7, 50, 103, 104], could usefully integrate these concepts and mechanisms. This would allow for direct comparisons with electrophysiological experimental data that may aid not only in data-based model identification but also reveal the fundamental theory of speech coding in the brain.

The practical implications of studies on neural oscillations and their ability to partly synchronize with external stimuli are important for the treatment of a variety of pathologies. For example, an experiment found a relationship between the synchronization of delta- and gamma-band networks and semantic fluency in post-stroke chronic aphasia [105]. Similarly, a study found tracking of theta rhythms but not delta rhythms in the logopenic variant of primary progressive aphasia, which may indicate ineffective top-down coding [106]. Dyslexia is another pathology that has been associated with disturbances in low-frequency rhythm tracking [4, 107–111]. According to the rise-time theory of dyslexia, reading difficulties result from the complexity of tracking the amplitude modulation of external signals, which leads to difficulties in speech perception, phonological processing, and, ultimately, reading [111]. For instance, children with dyslexia have abnormal delta rhythm phase alignment when interpreting rhythmic syllable sequences, which affects speech representation [112]. Likewise, children with dyslexia showed impaired tapping to a metronome beat with a frequency of 2 Hz [113]. Another study compared neural responses to speech and non-speech sounds in healthy people and those with dyslexia, revealing that healthy people had stronger delta-band responses in the right hemisphere and gamma-band responses in the left hemisphere [114]. In these experiments, difficulties in tracking low-frequency external rhythms were correlated with phonetic perception problems. It is noteworthy that individuals with dyslexia can compensate for their phonological perception deficits with semantic context, i.e., top-down compensatory mechanisms [115–117].

Taken together, these findings suggest that disturbances in low-frequency rhythm synchronization may be a factor that accompanies the progression of aphasia and dyslexia. Thus, the success of transcranial electrical stimulation in treating these conditions may be partially explained by the synchronization of neural oscillations with external stimuli [118–120]. Clearly, the practical application of such research must rely heavily on a solid theoretical framework capable of predicting treatment effects, developing hypotheses, and developing experimental and treatment protocols. The BRyBI model can provide such a theoretical basis.

In summary, our results shed light on the intrinsic constraints and compensatory mechanisms of human speech perception. At the same time, they offer a potential alternative and challenge to the prevalent AI NLP approaches to speech processing, pointing out how brain mechanisms may allow for robust speech recognition with high computational efficiency, even under conditions where LLMs appear to perform poorly.

## 4 Methods

Predictive coding implies two directions of information flow: top-down and bottom-up. Top-down flow is provided by constructing the generative model (GM) and passing predictions from higher abstract levels to the early sensory areas. Bottom-up flow propagates updates in predictions during the inference process provided by the DEM algorithm [70].

GM is essentially a stochastic dynamical system that has a hierarchical structure. Each level is formed by two types of variables: hidden and causal states. The hidden states are ruled by differential equations. The causal states serve to transfer information from the top to the bottom levels and represent predictions inferred from the internal model of the world. They are formed as nonlinear transformations of the hidden states. Thus, GM maintains a top-down information flow. The BRyBI model consists of two levels and is formalized as follows:

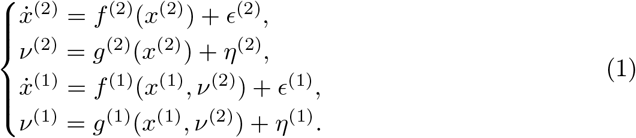

Here *x*^(*i*)^ is the hidden state of *i*-level with a noise *ϵ*^(*i*)^, *ν*^(*i*)^ is the corresponding causal state with a noise *η*^(*i*)^. The function *f* ^(*i*)^ determines a form of differential equations for the hidden state *x*^(*i*)^. The function *g*^(*i*)^ determines the nonlinear transformation of *x*^(*i*)^ taking into account information from the level above.

The bottom and top levels in BRyBI model mimic speech processing in the primary auditory cortex (pAC) and the associative auditory cortex (aAC), respectively (Fig. 1). On the top level, delta rhythm switches words. On the bottom level, coupled theta and gamma rhythms code acoustic signal of speech.

### 4.1 The top level

The top level is the level of context candidates that are alternating in delta rhythm. Here we define a delta-timescale:

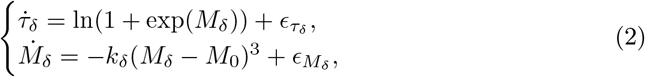

where the delta-timescale *τ*_*δ*_ is phase modulated by *M*_*δ*_. The parameter *k*_*δ*_ constrains the timescale in the delta band. *M*_*δ*_ potentially takes values from [*−∞, ∞*]. The function softplus(*M*_*δ*_) = ln(1 + exp(*M*_*δ*_)) maps the values to [0, +*∞*]. When *M*_*δ*_ = *M*_0_ the delta-wave does indeed have an average delta-rhythm frequency. In order to change this state, it is necessary to increase / decrease the noise 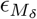;. Thus, the more the frequency differs from the delta rhythm, the more difficult it is to obtain the corresponding *M*_*δ*_ at the expense of noise (Fig. S1).

Delta-waves are constructed as follows:

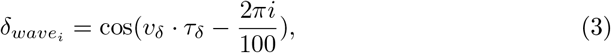

where 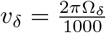, *Ω*_*δ*_ = 2.9 Hz is the average frequency of delta rhythm, and 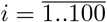. We designed delta waves in this way in order to simply generate a trigger in a certain phase of delta rhythm as *T*_*δ*_ = softmax_*i*_(*δ*_*waves*_). The delta trigger switches phrases by abruptly increasing from 0 to 1.

Phrases are chosen from the language randomly. The relative probability of each phrase accumulates in the variable **w**:

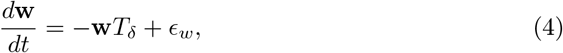

On this level, hidden states are (*τ*_*δ*_, *M*_*δ*_) and **w**. Causal states are 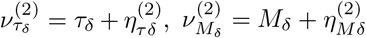 and word probabilities *ν*^(2)^ = softmax 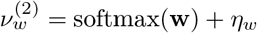.

### 4.2 The bottom level

The theta-timescale defines a window of syllable coding. We use the same model as for the delta-timescale:

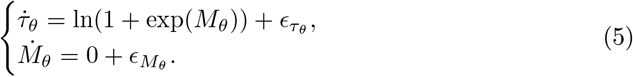

Following the example of previous similar models [23, 73, 98], the GM splits each syllable into 8 parts. It allows more flexibility in shaping the auditory spectrogram of syllables and phonemes. The gamma waves are constructed as a nonlinear function of *τ*_*θ*_ and each gamma wave has a period equal to 1*/*8 of the period of a syllable:

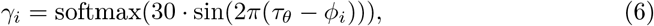

where *ϕ* = *i/*8, indexes *i* = 0..7.

Syllable selection is a crucial module in speech interpretation. In BRyBI, we want to enter the context of a certain phrase for its constitutive syllables. A phrase is basically an ordered sequence of syllables. In this definition, it is convenient to represent it as a matrix of syllables (Fig. 6):

**Fig. 6.**
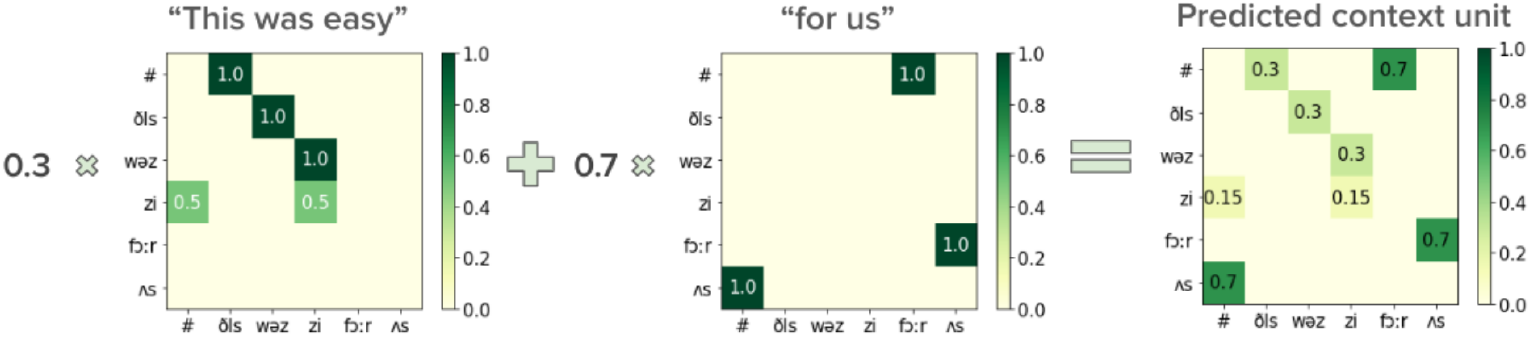
Example of context unit representation and construction as the sum of word matrices normalized by probabilities from the top level.

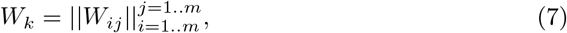

where *m* is a whole number of syllables in the language. An element *W*_*ij*_ = 1 if *j*-th syllable follows the *i*-th syllable; otherwise, *W*_*ij*_ = 0.

The context matrix is defined as the weighted sum of matrices of syllables: 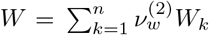, *n* is a number of words and phrases in the dictionary. If exactly one word were chosen in the variable **w**, i.e., only one value in the vector was equal to 1, and all the rest were equal to zero, then such a sum would choose from all matrices *W* only the one corresponding to the current word. The matrix *W* changes dynamically depending on the word/phrase probabilities at the top level. The resulting matrix *W* can be interpreted as a probability of transition between syllables according to a context unit.

Syllable transition frequency inside a word / phrase is in the theta-band, whereas word switching frequency is in the delta-band. Hidden states of syllables are determined as follows:

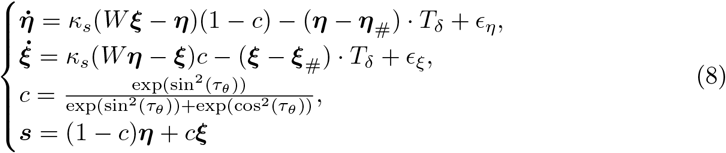

Here *η* and *ξ* are two syllable pointers. The equations determine a gradual transition from one syllable, which is pointed by *ξ*, to another syllable, which is pointed by *η*, with a speed *κ*_*s*_. These two pointers essentially follow their own theta wave. At the same time, the expected syllable from the context is encoded in half of the cycle; in the second half, it either occurs or it can knock out another syllable by error (if the context, for example, was chosen incorrectly).

On the bottom level, hidden states are the theta-timescale *τ*_*θ*_, theta modulation *M*_*θ*_, and syllable pointers *ξ* and *η*.

A theta-gamma code of an acoustic signal is generated as a convolution of syllable and gamma-units with predefined for each frequency band *f* tensors *P*_*fγθ*_:

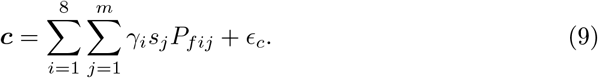

### 4.3 The dataset

To drive the model, we use both the spectrogram and syllable/prosodic envelopes (high-pass and low-pass envelopes, respectively) as inputs to the bottom level. The original dataset is a preprocessed TIMIT dataset [82]. Using the code from [23], we extracted the 6-channel spectra from the sentences. The extraction of syllable matrices is described in Section Methods: Dataset.

The extraction of syllable matrices is performed for each syllable in the sentence and consists of 3 steps:

1. A piece of the spectrum is extracted according to the boundaries of the syllable.
2. A piece of the spectrum is split into 8 equal segments.
3. Over time, each part is averaged. The result is eight 6-dimensional vectors, one for each scale.

The rhythm of speech is represented by phase modulation signals *M*_*θ*_ for theta and *M*_*θ*_ for delta rhythm. Then these modulation signals are directly compared with the generated signals.

### 4.4 Simulation details

Constructed GM is a likelihood in Bayesian inference framework. Since GM is a dynamical system, we run the inversion process by the DEM algorithm using the MATLAB library SPM12 [70]. Generated auditory spectrogram **c**, delta *M*_*δ*_ and theta *M*_*θ*_ modulations are compared with an input spectrogram, the prosodic envelope, and the syllabic envelope, respectively. DEM produces joint distributions for all hidden and causal variables, which are used in the model to recognize syllables and phrases. This process involves the inference of GM state trajectories as samples from Gaussian distributions. The covariance of these distributions for each state is different. Precisions (inverted covariance) for each state are shown in Table 1.

**Table 1.**
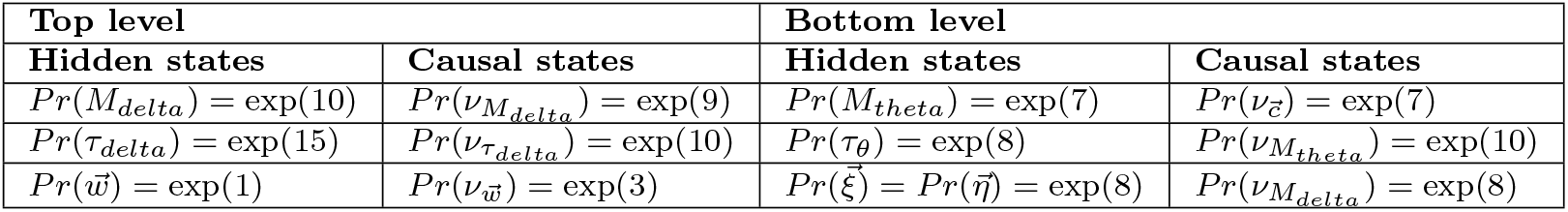
Precision values for model inversion by the DEM algorithm.

### 4.5 Analysis details

We calculated uncertainty as entropy: *x* = *−* ∑ *p*_*i*_ log *p*_*i*_. For all time steps in the data, we collected point pairs (*x, y*). Buckets were chosen on the interval (*−∞*; *∞*) in order to save the same number of points in each bucket. Table 2 shows uncertainty values included in each bucket.

**Table 2.** Uncertainty values are integrated into buckets.

| Number of bucket | Interval of uncertainty |
| --- | --- |
| 0 | $[-\text{inf}; 4.82\text{e-}03)$ |
| 1 | $[4.82\text{e-}03; 5.78\text{e-}03)$ |
| 2 | $[5.78\text{e-}03; 6.81\text{e-}03)$ |
| 3 | $[6.81\text{e-}03; 8.36\text{e-}03)$ |
| 4 | $[8.36\text{e-}03; 1.53\text{e-}02)$ |
| 5 | $[1.53\text{e-}02; 1.19\text{e-}01)$ |
| 6 | $[1.19\text{e-}01; 5.66\text{e-}01)$ |
| 7 | $[5.66\text{e-}01; \text{inf})$ |

Rhythm decoupling is calculated for delta and theta rhythm as a difference between predicted modulation signal and input modulation signal: (*M*_*predicted*_*−M*_*input*_)^2^, where *M* is a modulation of delta (*M*_*δ*_) or theta (*M*_*theta*_) rhythm.

## Acknowledgments

We thank Heike Stein for valuable comments on the manuscript. This publication is supported by the Brain Program of the IDEAS Research Center and the Vernadski scholarship. The research in part through computational resources of HPC facilities at HSE University. BSG was supported by CNRS, INSERM.

## Notes

### Competing Interest Statement

The authors have declared no competing interest.

### Summary of Updates

- Change the title; - Add an author; - Complete acknowledgements;

## References

[1] Viemeister, N.F., Wakefield, G.H.: Temporal integration and multiple looks. The Journal of the Acoustical Society of America 90(2), 858–865 (1991)

[2] Saberi, K., Perrott, D.R.: Cognitive restoration of reversed speech. Nature 398(6730), 760–760 (1999)

[3] Poeppel, D.: The analysis of speech in different temporal integration windows: cerebral lateralization as ‘asymmetric sampling in time’. Speech communication 41(1), 245–255 (2003)

[4] Ding, N., Melloni, L., Zhang, H., Tian, X., Poeppel, D.: Cortical tracking of hierarchical linguistic structures in connected speech. Nature neuroscience 19(1), 158–164 (2016)

[5] Ghitza, O.: Acoustic-driven delta rhythms as prosodic markers. Language, Cognition and Neuroscience 32(5), 545–561 (2017)

[6] Poeppel, D.: The maps problem and the mapping problem: two challenges for a cognitive neuroscience of speech and language. Cognitive neuropsychology 29(1-2), 34–55 (2012)

[7] Giraud, A.-L., Poeppel, D.: Cortical oscillations and speech processing: emerging computational principles and operations. Nature neuroscience 15(4), 511–517 (2012)

[8] Ronconi, L., Oosterhof, N.N., Bonmassar, C., Melcher, D.: Multiple oscillatory rhythms determine the temporal organization of perception. Proceedings of the National Academy of Sciences 114(51), 13435–13440 (2017)

[9] Chen, B., Ciria, L.F., Hu, C., Ivanov, P.C.: Ensemble of coupling forms and networks among brain rhythms as function of states and cognition. Communications Biology 5(1), 82 (2022)

[10] Buzsáki, G., Watson, B.O.: Brain rhythms and neural syntax: implications for efficient coding of cognitive content and neuropsychiatric disease. Dialogues in clinical neuroscience (2022)

[11] Hyafil, A., Giraud, A.-L., Fontolan, L., Gutkin, B.: Neural cross-frequency coupling: connecting architectures, mechanisms, and functions. Trends in neurosciences 38(11), 725–740 (2015)

[12] Gross, J., Hoogenboom, N., Thut, G., Schyns, P., Panzeri, S., Belin, P., Garrod, S.: Speech rhythms and multiplexed oscillatory sensory coding in the human brain. PLoS biology 11(12), 1001752 (2013)

[13] Teng, X., Tian, X., Rowland, J., Poeppel, D.: Concurrent temporal channels for auditory processing: Oscillatory neural entrainment reveals segregation of function at different scales. PLoS biology 15(11), 2000812 (2017)

[14] Teng, X., Tian, X., Doelling, K., Poeppel, D.: Theta band oscillations reflect more than entrainment: behavioral and neural evidence demonstrates an active chunking process. European Journal of Neuroscience 48(8), 2770–2782 (2018)

[15] Teng, X., Poeppel, D.: Theta and gamma bands encode acoustic dynamics over wide-ranging timescales. Cerebral cortex 30(4), 2600–2614 (2020)

[16] Aiken, S.J., Picton, T.W.: Human cortical responses to the speech envelope. Ear and hearing 29(2), 139–157 (2008)

[17] Luo, H., Poeppel, D.: Phase patterns of neuronal responses reliably discriminate speech in human auditory cortex. Neuron 54(6), 1001–1010 (2007)

[18] Doelling, K.B., Arnal, L.H., Ghitza, O., Poeppel, D.: Acoustic landmarks drive delta–theta oscillations to enable speech comprehension by facilitating perceptual parsing. Neuroimage 85, 761–768 (2014)

[19] Mesgarani, N., Cheung, C., Johnson, K., Chang, E.F.: Phonetic feature encoding in human superior temporal gyrus. Science 343(6174), 1006–1010 (2014)

[20] Tang, C., Hamilton, L., Chang, E.: Intonational speech prosody encoding in the human auditory cortex. Science 357(6353), 797–801 (2017)

[21] Murphy, E.: A theta-gamma neural code for feature set composition with phase-entrained delta nestings. UCL Work. Pap. Linguist 28, 1–23 (2016)

[22] Hyafil, A., Fontolan, L., Kabdebon, C., Gutkin, B., Giraud, A.-L.: Speech encoding by coupled cortical theta and gamma oscillations. Elife 4, 06213 (2015)

[23] Hovsepyan, S., Olasagasti, I., Giraud, A.-L.: Combining predictive coding and neural oscillations enables online syllable recognition in natural speech. Nature communications 11(1), 1–12 (2020)

[24] Ghitza, O., Greenberg, S.: On the possible role of brain rhythms in speech perception: intelligibility of time-compressed speech with periodic and aperiodic insertions of silence. Phonetica 66(1-2), 113–126 (2009)

[25] Cernak, M., Asaei, A., Hyafil, A.: Cognitive speech coding: examining the impact of cognitive speech processing on speech compression. IEEE Signal Processing Magazine 35(3), 97–109 (2018)

[26] Miller, G.A., Licklider, J.C.: The intelligibility of interrupted speech. The Journal of the Acoustical Society of America 22(2), 167–173 (1950)

[27] Huggins, A.: Temporally segmented speech. Perception & Psychophysics 18, 149–157 (1975)

[28] Garvey, W.D.: The intelligibility of speeded speech. Journal of experimental psychology 45(2), 102 (1953)

[29] Ghitza, O.: Behavioral evidence for the role of cortical θ oscillations in determining auditory channel capacity for speech. Frontiers in psychology 5, 652 (2014)

[30] Borges, A.F.T., Giraud, A.-L., Mansvelder, H.D., Linkenkaer-Hansen, K.: Scale-free amplitude modulation of neuronal oscillations tracks comprehension of accelerated speech. Journal of Neuroscience 38(3), 710–722 (2018)

[31] Giroud, J., Lerousseau, J.P., Pellegrino, F., Morillon, B.: The channel capacity of multilevel linguistic features constrains speech comprehension. Cognition 232, 105345 (2023)

[32] Penn, L.R., Ayasse, N.D., Wingfield, A., Ghitza, O.: The possible role of brain rhythms in perceiving fast speech: Evidence from adult aging. The Journal of the Acoustical Society of America 144(4), 2088–2094 (2018)

[33] Gransier, R., Peeters, S., Wouters, J.: The importance of temporal-fine structure to perceive time-compressed speech with and without the restoration of the syllabic rhythm. Scientific Reports 13(1), 2874 (2023)

[34] Mai, G., Peng, G.: Optimal syllabic rates and processing units in perceiving mandarin spoken sentences. In: INTERSPEECH, pp. 2477–2480 (2011)

[35] Stephenson, C., Feather, J., Padhy, S., Elibol, O., Tang, H., McDermott, J., Chung, S.: Untangling in invariant speech recognition. Advances in neural information processing systems 32 (2019)

[36] Greenberg, S., Kingsbury, B.E.: The modulation spectrogram: In pursuit of an invariant representation of speech. In: 1997 IEEE International Conference on Acoustics, Speech, and Signal Processing, vol. 3, pp. 1647–1650 (1997). IEEE

[37] Kösem, A., Bosker, H.R., Meyer, A.S., Jensen, O., Hagoort, P.: Neural entrainment reflects temporal predictions guiding speech comprehension. In: The Eighth Annual Meeting of the Society for the Neurobiology of Language (snl 2016) (2016)

[38] Kösem, A., Bosker, H.R., Takashima, A., Meyer, A., Jensen, O., Hagoort, P.: Neural entrainment determines the words we hear. Current Biology 28(18), 2867–2875 (2018)

[39] Okada, K., Rong, F., Venezia, J., Matchin, W., Hsieh, I.-H., Saberi, K., Serences, J.T., Hickok, G.: Hierarchical organization of human auditory cortex: evidence from acoustic invariance in the response to intelligible speech. Cerebral Cortex 20(10), 2486–2495 (2010)

[40] Evans, S., Davis, M.H.: Hierarchical organization of auditory and motor representations in speech perception: evidence from searchlight similarity analysis. Cerebral cortex 25(12), 4772–4788 (2015)

[41] Obleser, J., Leaver, A.M., VanMeter, J., Rauschecker, J.P.: Segregation of vowels and consonants in human auditory cortex: evidence for distributed hierarchical organization. Frontiers in psychology 1, 232 (2010)

[42] Heer, W.A., Huth, A.G., Griffiths, T.L., Gallant, J.L., Theunissen, F.E.: The hierarchical cortical organization of human speech processing. Journal of Neuroscience 37(27), 6539–6557 (2017)

[43] Caucheteux, C., Gramfort, A., King, J.-R.: Evidence of a predictive coding hierarchy in the human brain listening to speech. Nature Human Behaviour, 1–12 (2023)

[44] Longacre, R.E.: Hierarchy in language. Method and theory in linguistics, 173–195 (1970)

[45] Molinaro, N., Lizarazu, M., Baldin, V., Pérez-Navarro, J., Lallier, M., Ríos-López, P.: Speech-brain phase coupling is enhanced in low contextual semantic predictability conditions. Neuropsychologia 156, 107830 (2021)

[46] Ten Oever, S., Carta, S., Kaufeld, G., Martin, A.E.: Neural tracking of phrases in spoken language comprehension is automatic and task-dependent. Elife 11, 77468 (2022)

[47] Herbst, S.K., Obleser, J.: Implicit temporal predictability enhances pitch discrimination sensitivity and biases the phase of delta oscillations in auditory cortex. NeuroImage 203, 116198 (2019)

[48] Kaufeld, G., Bosker, H.R., Ten Oever, S., Alday, P.M., Meyer, A.S., Martin, A.E.: Linguistic structure and meaning organize neural oscillations into a content-specific hierarchy. Journal of Neuroscience 40(49), 9467–9475 (2020)

[49] Hannemann, R., Obleser, J., Eulitz, C.: Top-down knowledge supports the retrieval of lexical information from degraded speech. Brain research 1153, 134–143 (2007)

[50] Forseth, K.J., Hickok, G., Rollo, P., Tandon, N.: Language prediction mechanisms in human auditory cortex. Nature communications 11(1), 5240 (2020)

[51] Ding, R., Oever, S., Martin, A.E.: Pronoun resolution via reinstatement of referent-related activity in the delta band. bioRxiv, 2023–04 (2023)

[52] Barczak, A., O’Connell, M.N., McGinnis, T., Ross, D., Mowery, T., Falchier, A., Lakatos, P.: Top-down, contextual entrainment of neuronal oscillations in the auditory thalamocortical circuit. Proceedings of the National Academy of Sciences 115(32), 7605–7614 (2018)

[53] Ding, N., Chatterjee, M., Simon, J.Z.: Robust cortical entrainment to the speech envelope relies on the spectro-temporal fine structure. Neuroimage 88, 41–46 (2014)

[54] Myers, B.R., Lense, M.D., Gordon, R.L.: Pushing the envelope: Developments in neural entrainment to speech and the biological underpinnings of prosody perception. Brain sciences 9(3), 70 (2019)

[55] Molinaro, N., Lizarazu, M.: Delta (but not theta)-band cortical entrainment involves speech-specific processing. European Journal of Neuroscience 48(7), 2642–2650 (2018)

[56] Attaheri, A., Choisdealbha, Á.N., Di Liberto, G.M., Rocha, S., Brusini, P., Mead, N., Olawole-Scott, H., Boutris, P., Gibbon, S., Williams, I., et al.: Delta-and theta-band cortical tracking and phase-amplitude coupling to sung speech by infants. NeuroImage 247, 118698 (2022)

[57] Giroud, J., Trébuchon, A., Schön, D., Marquis, P., Liegeois-Chauvel, C., Poeppel, D., Morillon, B.: Asymmetric sampling in human auditory cortex reveals spectral processing hierarchy. PLoS biology 18(3), 3000207 (2020)

[58] Rimmele, J.M., Poeppel, D., Ghitza, O.: Acoustically driven cortical d oscillations underpin prosodic chunking. Eneuro 8(4) (2021)

[59] Roehm, D., Schlesewsky, M., Bornkessel, I., Frisch, S., Haider, H.: Fractionating language comprehension via frequency characteristics of the human eeg. Neuroreport 15(3), 409–412 (2004)

[60] Donhauser, P.W., Baillet, S.: Two distinct neural timescales for predictive speech processing. Neuron 105(2), 385–393 (2020)

[61] Bai, F., Meyer, A.S., Martin, A.E.: Neural dynamics differentially encode phrases and sentences during spoken language comprehension. PLoS Biology 20(7), 3001713 (2022)

[62] Park, H., Thut, G., Gross, J.: Predictive entrainment of natural speech through two fronto-motor top-down channels. Language, Cognition and Neuroscience 35(6), 739–751 (2020)

[63] Meyer, L., Henry, M.J., Gaston, P., Schmuck, N., Friederici, A.D.: Linguistic bias modulates interpretation of speech via neural delta-band oscillations. Cerebral Cortex 27(9), 4293–4302 (2017)

[64] Arnal, L.H., Doelling, K.B., Poeppel, D.: Delta–beta coupled oscillations underlie temporal prediction accuracy. Cerebral Cortex 25(9), 3077–3085 (2015)

[65] Doelling, K.B., Arnal, L.H., Assaneo, M.F.: Adaptive oscillators support bayesian prediction in temporal processing. PLOS Computational Biology 19(11), 1011669 (2023)

[66] Nabé, M., Schwartz, J.-L., Diard, J.: Cosmo-onset: A neurally-inspired computational model of spoken word recognition, combining top-down prediction and bottom-up detection of syllabic onsets. Frontiers in Systems Neuroscience, 75 (2021)

[67] Ghitza, O.: On the role of theta-driven syllabic parsing in decoding speech: intelligibility of speech with a manipulated modulation spectrum. Frontiers in psychology 3, 238 (2012)

[68] Rao, R.P., Ballard, D.H.: Predictive coding in the visual cortex: a functional interpretation of some extra-classical receptive-field effects. Nature neuroscience 2(1), 79–87 (1999)

[69] Bastos, A.M., Usrey, W.M., Adams, R.A., Mangun, G.R., Fries, P., Friston, K.J.: Canonical microcircuits for predictive coding. Neuron 76(4), 695–711 (2012)

[70] Friston, K., Kiebel, S.: Predictive coding under the free-energy principle. Philosophical transactions of the Royal Society B: Biological sciences 364(1521), 1211–1221 (2009)

[71] Heilbron, M., Chait, M.: Great expectations: is there evidence for predictive coding in auditory cortex? Neuroscience 389, 54–73 (2018)

[72] Peelle, J.E., Johnsrude, I., Davis, M.H.: Hierarchical processing for speech in human auditory cortex and beyond. Frontiers in human neuroscience 4, 51 (2010)

[73] Su, Y., MacGregor, L.J., Olasagasti, I., Giraud, A.-L.: A deep hierarchy of predictions enables online meaning extraction in a computational model of human speech comprehension. Plos Biology 21(3), 3002046 (2023)

[74] Friston, K.J., Sajid, N., Quiroga-Martinez, D.R., Parr, T., Price, C.J., Holmes, E.: Active listening. Hearing research 399, 107998 (2021)

[75] Friston, K.J., Parr, T., Yufik, Y., Sajid, N., Price, C.J., Holmes, E.: Generative models, linguistic communication and active inference. Neuroscience & Biobehavioral Reviews 118, 42–64 (2020)

[76] Zhao, B., Dang, J., Zhang, G., Unoki, M.: Cortical oscillatory hierarchy for natural sentence processing. In: INTERSPEECH, pp. 125–129 (2020)

[77] Poeppel, D., Idsardi, W.J., Van Wassenhove, V.: Speech perception at the interface of neurobiology and linguistics. Philosophical Transactions of the Royal Society B: Biological Sciences 363(1493), 1071–1086 (2008)

[78] Chandrasekaran, C., Trubanova, A., Stillittano, S., Caplier, A., Ghazanfar, A.A.: The natural statistics of audiovisual speech. PLoS computational biology 5(7), 1000436 (2009)

[79] Ghitza, O.: Linking speech perception and neurophysiology: speech decoding guided by cascaded oscillators locked to the input rhythm. Frontiers in psychology 2, 130 (2011)

[80] Rumelhart, D.E., McClelland, J.L.: An interactive activation model of context effects in letter perception: Ii. the contextual enhancement effect and some tests and extensions of the model. Psychological review 89(1), 60 (1982)

[81] McClelland, J.L., Rumelhart, D.E.: An interactive activation model of context effects in letter perception: I. an account of basic findings. Psychological review 88(5), 375 (1981)

[82] Garofolo, J.S.: Timit acoustic phonetic continuous speech corpus. Linguistic Data Consortium, 1993 (1993)

[83] Baevski, A., Zhou, Y., Mohamed, A., Auli, M.: wav2vec 2.0: A framework for self-supervised learning of speech representations. Advances in neural information processing systems 33, 12449–12460 (2020)

[84] Tezcan, F., Weissbart, H., Martin, A.E.: A tradeoff between acoustic and linguistic feature encoding in spoken language comprehension. Elife 12, 82386 (2023)

[85] Radford, A., Kim, J.W., Xu, T., Brockman, G., McLeavey, C., Sutskever, I.: Robust speech recognition via large-scale weak supervision. In: International Conference on Machine Learning, pp. 28492–28518 (2023). PMLR

[86] MS Azure, Speech to Text. https://azure.microsoft.com/en-us/products/ai-services/speech-to-text Accessed 2024-04-23

[87] Google, Speech to Text. https://cloud.google.com/speech-to-text Accessed 2024-04-23

[88] Schrimpf, M., Blank, I.A., Tuckute, G., Kauf, C., Hosseini, E.A., Kanwisher, N., Tenenbaum, J.B., Fedorenko, E.: The neural architecture of language: Integrative modeling converges on predictive processing. Proceedings of the National Academy of Sciences 118(45), 2105646118 (2021)

[89] Mehrish, A., Majumder, N., Bharadwaj, R., Mihalcea, R., Poria, S.: A review of deep learning techniques for speech processing. Information Fusion, 101869 (2023)

[90] Martin, A.E.: A compositional neural architecture for language. Journal of Cognitive Neuroscience 32(8), 1407–1427 (2020)

[91] Slaats, S., Weissbart, H., Schoffelen, J.-M., Meyer, A.S., Martin, A.E.: Delta-band neural responses to individual words are modulated by sentence processing. Journal of Neuroscience 43(26), 4867–4883 (2023)

[92] Lau, E.F., Phillips, C., Poeppel, D.: A cortical network for semantics:(de) constructing the n400. Nature reviews neuroscience 9(12), 920–933 (2008)

[93] Ten Oever, S., Schroeder, C.E., Poeppel, D., Van Atteveldt, N., Zion-Golumbic, E.: Rhythmicity and cross-modal temporal cues facilitate detection. Neuropsychologia 63, 43–50 (2014)

[94] Rimmele, J.M., Morillon, B., Poeppel, D., Arnal, L.H.: Proactive sensing of periodic and aperiodic auditory patterns. Trends in cognitive sciences 22(10), 870–882 (2018)

[95] Etard, O., Reichenbach, T.: Neural speech tracking in the theta and in the delta frequency band differentially encode clarity and comprehension of speech in noise. Journal of Neuroscience 39(29), 5750–5759 (2019)

[96] Canales-Johnson, A., Borges, A.F.T., Komatsu, M., Fujii, N., Fahrenfort, J.J., Miller, K.J., Noreika, V.: Broadband dynamics rather than frequency-specific rhythms underlie prediction error in the primate auditory cortex. Journal of Neuroscience 41(45), 9374–9391 (2021)

[97] Giraud, A.-L.: Oscillations for all^−^\_ () _/^−^? a commentary on meyer, sun & martin (2020). Language, Cognition and Neuroscience 35(9), 1106–1113 (2020)

[98] Hovsepyan, S., Olasagasti, I., Giraud, A.-L.: Rhythmic modulation of prediction errors: A top-down gating role for the beta-range in speech processing. PLOS Computational Biology 19(11), 1011595 (2023)

[99] Hummel, J.E., Holyoak, K.J.: Distributed representations of structure: A theory of analogical access and mapping. Psychological review 104(3), 427 (1997)

[100] Doumas, L.A., Hummel, J.E., Sandhofer, C.M.: A theory of the discovery and predication of relational concepts. Psychological review 115(1), 1 (2008)

[101] Shastri, L.: Types and quantifiers in shruti–a connectionist model of rapid reasoning and relational processing. In: International Workshop on Hybrid Neural Systems, pp. 28–45 (1998). Springer

[102] Martin, A.E., Doumas, L.A.: A mechanism for the cortical computation of hierarchical linguistic structure. PLoS biology 15(3), 2000663 (2017)

[103] Poeppel, D., Assaneo, M.F.: Speech rhythms and their neural foundations. Nature reviews neuroscience 21(6), 322–334 (2020)

[104] Stanley, D.A., Falchier, A.Y., Pittman-Polletta, B.R., Lakatos, P., Whittington, M.A., Schroeder, C.E., Kopell, N.J.: Flexible reset and entrainment of delta oscillations in primate primary auditory cortex: modeling and experiment. BioRxiv, 812024 (2019)

[105] Mehraram, R., Kries, J., De Clercq, P., Vandermosten, M., Francart, T.: Eeg reveals brain network alterations in chronic aphasia during natural speech listening. bioRxiv, 2023–03 (2023)

[106] Dial, H.R., Gnanateja, G.N., Tessmer, R.S., Gorno-Tempini, M.L., Chandrasekaran, B., Henry, M.L.: Cortical tracking of the speech envelope in logopenic variant primary progressive aphasia. Frontiers in human neuroscience 14, 597694 (2021)

[107] Lallier, M., Lizarazu, M., Molinaro, N., Bourguignon, M., Ríos-López, P., Carreiras, M.: From auditory rhythm processing to grapheme-to-phoneme conversion: How neural oscillations can shed light on developmental dyslexia. Reading and Dyslexia: From Basic Functions to Higher Order Cognition, 147–163 (2018)

[108] Di Liberto, G.M., Peter, V., Kalashnikova, M., Goswami, U., Burnham, D., Lalor, E.C.: Atypical cortical entrainment to speech in the right hemisphere underpins phonemic deficits in dyslexia. NeuroImage 175, 70–79 (2018)

[109] Power, A.J., Colling, L.J., Mead, N., Barnes, L., Goswami, U.: Neural encoding of the speech envelope by children with developmental dyslexia. Brain and Language 160, 1–10 (2016)

[110] Destoky, F., Bertels, J., Niesen, M., Wens, V., Vander Ghinst, M., Rovai, A., Trotta, N., Lallier, M., De Tiège, X., Bourguignon, M.: The role of reading experience in atypical cortical tracking of speech and speech-in-noise in dyslexia. NeuroImage 253, 119061 (2022)

[111] Goswami, U.: Sensory theories of developmental dyslexia: three challenges for research. Nature Reviews Neuroscience 16(1), 43–54 (2015)

[112] Power, A.J., Mead, N., Barnes, L., Goswami, U.: Neural entrainment to rhythmic speech in children with developmental dyslexia. Frontiers in human neuroscience 7, 777 (2013)

[113] Thomson, J.M., Goswami, U.: Rhythmic processing in children with developmental dyslexia: auditory and motor rhythms link to reading and spelling. Journal of Physiology-Paris 102(1-3), 120–129 (2008)

[114] Lizarazu, M., Covella, L.S., Wassenhove, V., Rivière, D., Mizzi, R., Lehongre, K., Hertz-Pannier, L., Ramus, F.: Neural entrainment to speech and nonspeech in dyslexia: conceptual replication and extension of previous investigations. Cortex 137, 160–178 (2021)

[115] Klimovich-Gray, A., Di Liberto, G., Amoruso, L., Barrena, A., Agirre, E., Molinaro, N.: Increased top-down semantic processing in natural speech linked to better reading in dyslexia. NeuroImage 273, 120072 (2023)

[116] Giraud, A.-L., Ramus, F.: Neurogenetics and auditory processing in developmental dyslexia. Current opinion in neurobiology 23(1), 37–42 (2013)

[117] Lehongre, K., Morillon, B., Giraud, A.-L., Ramus, F.: Impaired auditory sampling in dyslexia: further evidence from combined fmri and eeg. Frontiers in human neuroscience 7, 454 (2013)

[118] Elsner, B., Kugler, J., Pohl, M., Mehrholz, J.: Transcranial direct current stimulation (tdcs) for improving aphasia in adults with aphasia after stroke. Cochrane Database of Systematic Reviews (5) (2019)

[119] Biou, E., Cassoudesalle, H., Cogné, M., Sibon, I., De Gabory, I., Dehail, P., Aupy, J., Glize, B.: Transcranial direct current stimulation in post-stroke aphasia rehabilitation: A systematic review. Annals of physical and rehabilitation medicine 62(2), 104–121 (2019)

[120] Xie, X., Hu, P., Tian, Y., Wang, K., Bai, T.: Transcranial alternating current stimulation enhances speech comprehension in chronic post-stroke aphasia patients: A single-blind sham-controlled study. Brain Stimulation: Basic, Translational, and Clinical Research in Neuromodulation 15(6), 1538–1540 (2022)

