## Supplementary for "A brain-rhythm based computational framework for semantic context and acoustic signal integration in speech processing" (from another version of this preprint: version 3, 2024-05-03)

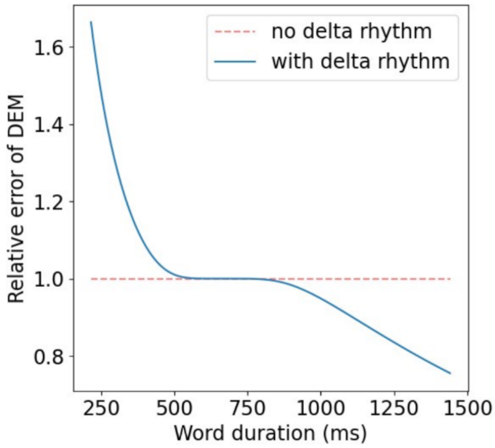


**Fig. 1** Noise sizing for delta rhythm constraint. For the equations of delta oscillations, we chose the functional form of the dependence and the noise such that, when adding the maximum noise, the oscillation frequency did not exceed the delta rhythm range.

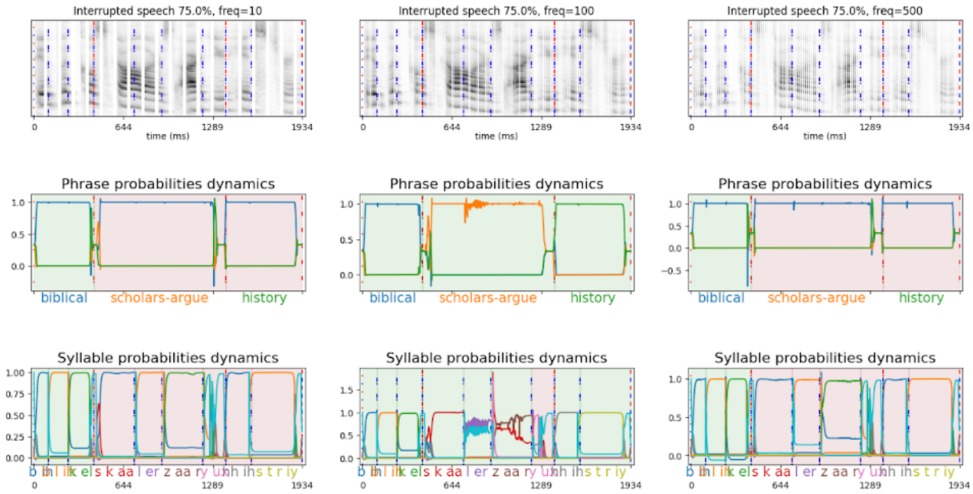


**Fig. 2** BRyBI performance for the "interrupted speech" experiment. As in the behavior experiments, BRyBI shows a drop in the articulation score in performance at the interruption frequency of 1 Hz that is caused by a long information loss interval (left). We also seepeaks in performance between 10 and 100 Hz, the optimal interruption rate allowing for recovery of missed information from context (middle). Finally, the drop at the interruption frequency $> 100$ Hz results from an impairment in phoneme decoding from a distorted spectrogram (right).

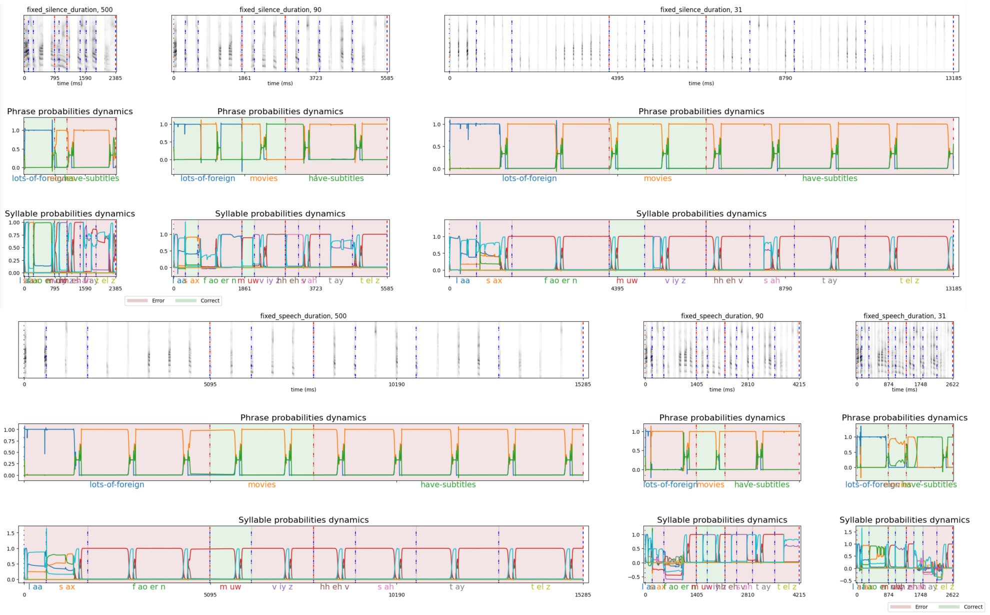


**Fig. 3** BRyBI performance for the temporally segmented speech experiment. Maintaining silent intervals at a consistent 200 msec while varying speech intervals reveals a decline in intelligibility for speech intervals ranging from 200 to 31 msec (Fig. 3G, blue line). Similarly, when speech-interval duration is fixed at approximately 63 msec, increasing silent-interval duration from 63 to 500 msec results in a decrease in intelligibility (Fig. 3G, orange line).

| Factor of compression | 1 | 1.25 | 1.5 | 1.65 | 1.75 | 1.875 | 2 | 2.5 | 3 |
|---|---|---|---|---|---|---|---|---|---|
| mean duration of syllables, ms | 199.3 | 159.4 | 132.9 | 122.6 | 113.9 | 106.3 | 99.7 | 79.7 | 66.5 |
| std, duration of syllables, ms | 105.3 | 84.2 | 70.2 | 64.8 | 60.2 | 56.2 | 52.6 | 42.1 | 35.1 |
| mean, duration of phrase, ms | 560.2 | 448.1 | 373.5 | 344.7 | 320.1 | 298.8 | 280.2 | 224.2 | 186.9 |
| std, duration of phrase, ms | 162.4 | 129.9 | 108.3 | 99.9 | 92.8 | 86.6 | 81.2 | 65.0 | 54.2 |

**Table 1** Statistics of syllable and prosodic phrase length under different factors of compression (in msec). Colors indicate the frequency range. Green: syllable frequency in 3-8 Hz and phrase frequency in 1-3 Hz, Orange: syllable frequency in 8-10 Hz and phrase frequency in 3-4 Hz, Red: syllable frequency is bigger than 10 Hz and phrase frequency is bigger than 4 Hz.
